# Signals of anticipation of reward and of mean reward rates in the human brain

**DOI:** 10.1101/754663

**Authors:** Roberto Viviani, Lisa Dommes, Julia Bosch, Michael Steffens, Anna Paul, Katharina L. Schneider, Julia C. Stingl, Petra Beschoner

**Author notes:** Corresponding author: Roberto Viviani, Institute of Psychology, University of Innsbruck, Innrain 52, 6020 Innsbruck, Austria, Psychiatry and Psychotherapy III, University of Ulm, Leimgrubenweg 12, 89075 Ulm, Germany.

## Abstract

Theoretical models of dopamine function stemming from reinforcement learning theory have emphasized the importance of prediction errors, which signal changes in the expectation of impending rewards. Much less is known about the effects of mean reward rates, which may be of motivational significance due to their role in computing the optimal effort put into exploiting reward opportunities. Here, we used a reinforcement learning model to design three functional neuroimaging studies and disentangle the effects of changes in reward expectations and mean reward rates, showing recruitment of specific regions in the brainstem regardless of prediction errors. While changes in reward expectations activated ventral striatal areas as in previous studies, mean reward rates preferentially modulated the substantia nigra/ventral tegmental area, deep layers of the superior colliculi, and a posterior pontomesencephalic region. These brainstem structures may work together to set motivation and attentional efforts levels according to perceived reward opportunities.

## 1 Introduction

The issue of the organization of motivational systems carries important implications in psychopathology. Examples are psychopathological states involving ‘lack of drive’ or lack of motivation/anhedonia (Pizzagalli et al. 2005; Nestler and Carlezon 2006; Treadway and Zald 2011; Der-Avakian and Markou 2012; Huys et al. 2013), or the difficulty to maintain sustained efforts to achieve goals, as in burn-out or neurasthenia (Maslach et al. 2001; Eriksen and Ursin 2002; Abbey and Garfinkel 1991). Motivational factors may also affect the propensity to sustain demanding cognitive efforts (Bromberg-Martin et al. 2010; Krebs et al. 2012; Kurzban et al. 2013). The identification of the processes regulating investment of efforts in the pursuit of goals may therefore contribute to mapping the candidate motivational components affected in important psychopathological states (NIMH RDoC Working Group 2011).

Studies in laboratory animals have provided abundant evidence of the relevance of appetitive reinforcers, or rewards, in motivation. One source of evidence goes back to early behavioural studies in laboratory animals on the effects of different amounts of reward on energizing behaviour (Crespi 1944). These findings provide the most direct behavioural evidence for the effect of reward rates on efforts invested in obtaining rewards. Subsequent studies showed that pharmacological manipulations of the dopamine system also lead to apparent energizing behavioural effects (Robbins and Everitt 1992; Berridge 2007; Robbins and Everitt 2007; Salamone and Correa 2012). These findings suggest that the dopamine system may be a key component of the motivational mechanism that links available reward rates with behavioural activation.

An influential model linking reward rates with motivational levels is based on the observation that, when averaged over time, available rewards provide an estimate of the ‘opportunity cost’ incurred in failing to undertake an action to secure the reward (Niv et al. 2005, 2007). Using a sophisticated version of reinforcement learning modelling (Sutton and Barto 1998), Niv et al. (2005) showed that, when the mean reward rate of actions is high, it makes sense to expend efforts to carry out the actions. This provides a normative justification for the existence of a mechanism that links estimates of mean available reward rates to motivational levels. However, several aspects of this proposed mechanism have yet to be clarified.

One such aspect is the relationship between the putative motivational process associated with different rates of reward, considered in the Niv et al. (2005) model, and the much more widely discussed form of motivation that is triggered by conditioned appetitive cues, which signal the availability of rewards (Estes 1943; Rescorla and Solomon 1967; Bolles 1972; Bindra 1974; Weingarten 1983; Berridge 2001). We will refer to this type of cue-bound behavioural activation as ‘incentive motivation’ (e.g., Bolles 1972). Considerable evidence involves dopamine signalling in incentive motivation through its double role in motivating responses and learning to predict rewards (Schultz 1998; Berridge and Robinson 1998; Ikemoto and Panksepp 1999; Dickinson et al. 2000; Alcaro et al. 2007; Salamone and Correa 2012). An important advance in understanding the effects of appetitive cues was made when it was realized that a wide range of neurophysiological data on dopamine discharge could be explained as the generation of a ‘prediction error’ as in computational models of reinforcement learning (Montague et al. 1996; Schultz et al. 1997; for recent reviews, see Schultz 2015; Daw and Tobler 2014). The dopamine signal is not generated by reward or conditioned cues per se, but only by the unexpected appearance of reward or of cues that predict reward (Schultz 1998). That is, the appearance of cues activates the dopamine system only when they provide new information on the upcoming reward. In this, the dopamine signal closely follows the prediction error signal of computational theories of learning (Waelti et al. 2001; Bayer and Glimcher 2005). Based on this computational model, an account of motivation on the basis of prediction error has been suggested, defining incentive motivation in terms of cues that change the current level of expectation of future rewards (Montague et al. 1996; Montague and Berns 2002; McClure et al. 2003a). Consistently with these theories, functional MRI studies in man have shown that informative cues, which change expectations on upcoming rewards and where the model generates a prediction error signal, are much more effective in activating areas corresponding to dopaminergic networks than non-informative cues, where the model generates no such signal (for example, cues made predictable by regularly alternating high and low reward conditions: Berns et al. 2001).

More recently, however, it has been increasingly clear that dopamine has many functions, and that not all of them may be modelled by the prediction error (Schultz 2015, 2016). Specifically, we are here concerned with the issue of whether prediction error offers a comprehensive model of motivation arising from different reward rates. Note, in particular, that despite the involvement of reward in both, the quantities representing detection of cues that inform on new reward opportunities and those representing probable rates of reward are computationally distinct. For example, when the reward contingencies are fully predicted, the cues convey no new information and no dopamine signal is observed (Montague et al. 1996). Even then, however, probable rates of reward would provide a useful estimate of opportunity costs. Indeed, it seems reasonable to assume from a normative point of view that acting to exploit opportunities is most appropriate when predictions about these opportunities are correct, and no prediction error is generated. This situation is known as the ‘consistency condition’ in the reinforcement learning literature (Sutton and Barto 1998). The consistency condition occurs when the temporal difference algorithm has learned to predict the subsequent rewards correctly. When this is the case, expected and obtained rewards match, and no changes in expectations take place. However, the algorithm internally keeps track of the expected reward rates.

In colloquial terms, this issue may be expressed as follows: incentive motivation may be triggered at the moment when we discover that a reward is potentially available, and the expectation of obtaining a reward is raised (prediction error). The motivational effect of such changes of expectation may consist, for example, in a focus on the reward associated with cue, and the activation of approach or seeking behaviour (Bindra 1974; Ikemoto and Panksepp 1999; Alcaro et al. 2007). In contrast, the motivation associated with different reward rates may be raised regardless of changes in expectations. An opportunity cost is incurred if we opt not to work to obtain that reward, even if our information about the availability of reward remains the same. If this were not the case, there would be no motivation to act in a context in which constant correct predictions of high reward rates signal high perceived opportunities. While in ordinary circumstances unexpected appetitive cues and reward rates may occur simultaneously or in complicated combinations, it may be informative to look at the neural substrates activated in these two conditions separately to address the issue of whether motivation from reward rates may be identified in terms of its associated neural substrates.

Several neurophysiological studies in laboratory animals provide data that suggest the importance of signals unrelated to prediction error. For example, a dopaminergic signal that could not be attributed to prediction error has been reported to occur when stimuli are presented in a ‘rewarding context’, i.e. when they are likely to be conducive to rewards (Kobayashi and Schultz 2014). Other midbrain signals associated with reward but that are not attributable to prediction error have been reported in the deep layers of the superior colliculi and in the peduncolopontine nuclei (Floresco et al. 2003; Redgrave and Gurney 2006). These neurophysiological data suggest that a neural substrate associated with opportunity costs, at the net of response rates or levels of effort, would be of importance to assess appetitive motivational mechanisms active in the early consummatory phases of the interaction with rewards.

Here, we carried out three functional MRI experiments that provided evidence for the activation of brainstem areas when disentangling the effects of prediction errors and mean reward rates. In all experiments a signal tracked expectations of mean reward rates in the brainstem was detected in the ventral tegmental area and the substantia nigra (VTA/SN). In contrast, the signal attributable to prediction error in the ventral striatum was affected by manipulations altering the predictability of the cue as predicted by the model.

## 2 Materials and Methods

### 2.1 Methods overview and rationale

To verify the hypothesis of a reward-related signal not associated with prediction error, we carried out three functional MRI experiments where levels of reward varied while their possible different effects on prediction error and reward opportunities may be identified. In contrast with previous studies, response levels were kept constant.

All experiments used variations of a basic task. Participants viewed a cue announcing the level of reward of a subsequent foraging patch, which could be either 1 or 20 cents at each correct response. In the foraging patch, a rapid sequence of target points appeared at the left or the right of the fixation point at fixed positions at slightly irregular intervals. Participants were required to press the left or the right button for targets appearing on the left or on the right, respectively (Figure 1A). In the foraging patches, each correct response always delivered either 1 or 20 cents, as announced by the preceding cues. All participants practiced the appropriate version of the task for 4 min. prior to the scan to establish the associations between cues and reward levels. In the scanner, high and low reward trials appeared in random order so as to make the cues informative about the forthcoming rewards.

**Figure 1.**
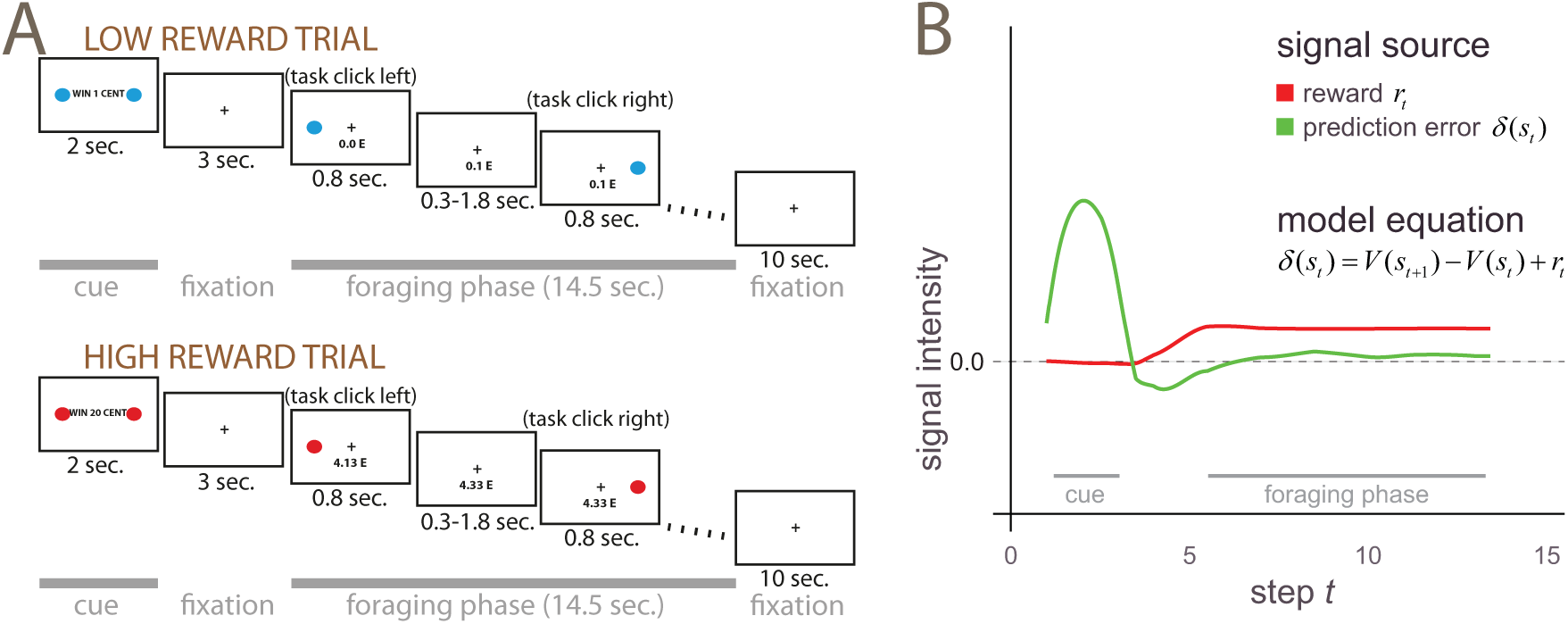
A: Schematic illustration of a low and a high reward trial in the task. After presentation of a cue, a rapid sequence of target episodes was presented rapidly during the foraging patch. B: Computer simulation of the two signal sources in the temporal difference model investigated in the paper (see the Appendix for details). The figure shows time-averaged signals generated by the algorithm after training. The prediction error signal *δ*, in green in the figure, represents changes in expected rewards, while *r* is the amount of reward effectively available in the foraging patch (in red). Smoothed over time, *r* gives the reward rate of the foraging patch. According to the model, the green curve corresponds to the incentive motivation signal, and the red curve to reward rates/perceived opportunities.

The design of this experiment was informed by the simplest reinforcement learning model (Sutton and Barto 1998) that captured the distinction between prediction errors and reward opportunity levels. This model consisted of the events of the cue and a series of events at the presentation of the targets. After learning, this model produces a prediction error signal at the presentation of the cue related to this change in expectations (Montague et al. 1996). The green curve in Figure 1B shows the course of the prediction error (the term *δ*(*s*_*t*_) of the equation shown in the Figure) from computer simulations of the model, after smoothing to better reflect the characteristics of the BOLD signal. Because no reward is provided at the presentation of the cue, reward rates remain at zero (mean reward rates are shown as a red curve in Figure 1B, corresponding to a temporally averaged version of the term *r*_*t*_ of the model equation). In contrast, during the foraging patches the dominant signal reflects mean levels of reward *r*_*t*_. There are no net changes in expectations of reward during the foraging patches, as the expectations informed by the preceding cue require no revision during this time. In this part of the trial the prediction error hovers around zero, and the consistency condition holds (more details on the temporal difference model, the consistency condition, and alternative modelling approaches are section 2.2. of the Methods). Note, however, that the targets resolved uncertainty about the side where they would appear. Hence, the task of detecting the targets and responding requires recruitment of processes to handle the uncertainty on the instantiation of targets. Therefore, an alternative interpretation of the qualitative difference between cues and foraging patches is that they provide information to resolve two types of uncertainty: about the amount of reward and about when and where targets appear.

A key aspect of the theoretical proposal of Niv et al. (2005) is that it addressed behaviour in the free operant setting, i.e. the setting in which the animal itself sets the pace of its efforts in responding to cues signalling possible rewards. In this setting, the energizing effects of cues and reward rates are directly observable as changes in response rates, determined jointly by the available rewards and the efforts required to obtain them. These changes are also consistent with the observed energizing effects of pharmacological manipulations of the dopamine system, noted above (Robbins and Everitt 1992; Berridge 2007; Robbins and Everitt 2007; Salamone and Correa 2012). Accordingly, to target these energizing effects previous work has focussed on the physical efforts prompted by rewards (Pessiglione et al. 2007), their effect on vigor of responding (Kurniawan et al. 2010; Guitart-Masip et al. 2011), or the estimation of cost-benefit trade-offs at different reward rates (Walton et al. 2006; Phillips et al. 2007). However, we were here interested in the neural substrates of representations of reward rates as such, i.e. at the net of estimates of effort costs or consequences on rates of responding. It seems reasonable to assume that, even if the costs of executing responses and the rate of responses are fixed, the opportunity cost signal used to regulate expenditure of efforts may still be computed. If this reasoning is correct, then we should observe activation of a neural substrate in association with reward rates when there is no prediction error and response costs and response rates are matched.

To match motor output between the low and high reward conditions, the awarded money was introduced as a virtual currency to obtain a movie theatre coupon at the end of the session. Participants would receive this coupon only if at the end of the task they had collected enough cents to equal the coupon value, and nothing otherwise. Participants were informed of the fact that they could not afford to miss more than few targets (including the 1 cent targets) during the experiment to reach the coupon price threshold. The sequence of trials was designed so that this threshold could be reached only towards the end of the task, enticing participants to keep working at the task up to the very last few targets of the last foraging block. The targets disappeared rapidly if participants did not respond immediately, forcing them to apply equal efforts over the whole task. For a real-life analogy, consider the activation of the dopamine system by fictitious rewards in a video game (Koepp et al. 1998).

Another desirable consequence of the introduction of a virtual currency was that from the standpoint of goal-directed control or of effortful, explicit executive processes there was no difference between low and high reward trials. The information conveyed by the different cues was essentially irrelevant: because also the 1 cent targets were needed to reach the threshold, to win the coupon participants just had to click on the correct side when targets appeared. However, there is ample evidence on the extent of processing of reward information in the brain even when this information is not required by the task (Bayer and Glimcher 2005). Bromberg-Martin and Hikosaka (2009), in particular, discuss “the puzzling fact that the brain devotes a great deal of neural effort to processing reward information even when this is not required to perform the task at hand” (p. 122 and citations therein). It has also been noted that “the dopamine system continued to compute the reward prediction error even when the behavioral policy of the animal was only weakly influenced by this computation.”

In the first experiment, we used the different characteristics of the signal at the cue and in the foraging patches in the temporal difference model to construct two contrasts on the data separating the effects of reward in changes in expectations and mean reward rates. Specifically, according to the model contrasts between the different trials types at the cue events detect differences in changes in reward expectations associated with prediction errors, while contrasts at the foraging patches detect differences in mean reward rates. Our purpose was to identify the neural substrates associated with informative cues and with different mean reward rates within the same experiment.

In the second and third experiments we manipulated the unpredictability of the cues and of the targets, respectively. The predictability of the cues was obtained by alternating high and low reward trials regularly (Berns et al. 2001), after informing participants of this fact. The predictability of targets was obtained by presenting them at the same regular intervals. According to the temporal difference model, these changes in predictability affect the signal conveyed by the prediction error, but not the signal tracking mean reward rates. The different sensitivity of these two signal sources to predictability therefore provides an additional opportunity to verify that the signals observed in the brain match the different computational properties described by the model.

### 2.2 Modelling

#### 2.2.1 Temporal difference models of the consistency condition

In its simplest version, the temporal difference algorithm (TD) models the process through which the ‘values’ of the states that may occur are learned. State values are an estimate of the expected rewards that can be earned in the future from that state, taking into account a temporal discounting term. ‘States’ are in this context a formalization of the steps through which the learner goes through when earning rewards. Formally, if we denote the state and reward at time step *t* as *s*_*t*_ and *r*_*t*_, the state value estimation function *V*(*s*_*t*_) may be written

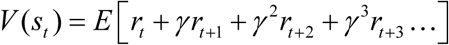

where *E* denotes expectation, and 0 < *γ* < 1 is the temporal discounting term. Here, we used an undiscounted version of the algorithm (*γ* = 1), known as TD(0) (Sutton and Barto 1998), since even in its simplest form the algorithm captures the distinction between prediction error and rewards matching expectations.

To learn state values from experience, the value of each state *V*(*s*_*t*_) where *t* indexes the sequence of the progression of the trial, is updated at each training step by incrementing it by a fraction of the prediction error *δ*(*s*_*t*_), i.e. the discrepancy between estimated and observed rewards in successive states:

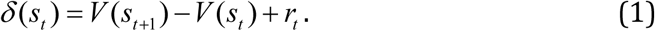

When rewards match expectations there is no prediction error. Setting the prediction error term *δ*(*s*_*t*_) to zero in equation 1, one obtains an equation that links obtained rewards to internal representations of value, known in the learning theory literature as the ‘consistency condition’ (Sutton and Barto 1998):

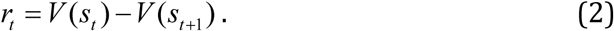

In the foraging patches, reward rates are equal to the differential valuation term *V*(*s*_*t*_) – *V*(*s*_*t* +1_) of this equation, which reflects the way in which internal estimates of expected reward are coded in this algorithm. Hence, in this phase of the task the constant reward rates of the model may just as well be viewed in terms of values internally assigned to the current and the next states, i.e. the perceived opportunities for reward at these states. We draw attention to this aspect of the model in the setting of our task, because constant reward rates in equilibrium with expectations may be better thought of as internal representations of opportunities than the pleasurable experience of reward itself. The experience of reward does not require work, and may be associated with hedonic processes that have been shown to be distinct from incentive motivation and dopamine function (Berridge 2007; Salamone and Correa 2012; Berridge and Kringelbach 2015). The fact that participants here received the reward only at the end of the task also reflects this view of what perceived opportunities ought to be. Note also that *V*(*s*_*t*_) – *V*(*s*_*t* +1_) refers to an expectation of reward that may come next, in contrast to *V*(*s*_*t*_).

Another perspective on the differential term *V*(*s*_*t*_) – *V*(*s*_*t* +1_) in the foraging patches, where values of states match expectations, is given by considering a more complex model explicitly representing the value of responses (referred to as *Q*-values in reinforcement learning theory). To choose the optimal response, the algorithm aims to learn the value of state-action pairs *Q*(*s*_*t*_, *a*_*t*_), instead of simply the value of states *V*(*s*_*t*_) (Sutton and Barto 1998). The values of state-action pairs is the value accruing from choosing the action *a*_*t*_ at the step *t* and following the optimal policy from the given step (the best choice of actions und the current estimate). Given that the optimal policy here is to always click when a target appears, the consistency condition equation for *r*_*t*_ when rewards match expectations becomes

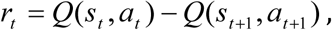

showing that the *r*_*t*_ term of the model is equivalent to the value of clicking relative to the value of inaction at the present episode, estimated from the long-term value of the states that follow under the optimal policy. In terms of internal estimates, differences in *r*_*t*_ in the foraging patches may represent information on the level of support present in the environment for the response associated with the collection of rewards. In conclusion, when rewards match expectations the term *r*_*t*_ takes similar roles in different versions of learning algorithms within the TD framework, but each of these versions draw attention on slightly different aspects of the information it may represent.

There are several aspects of the task in the foraging patch that our simple model fails to make explicit. One is that dopamine signalling may be sensitive to the variability in the timing of target appearance. The temporal difference model responds to a delay of the target by producing a negative prediction error. Conversely, anticipated targets produce a positive spike. Over time, the negative and positive spikes average out, so that the average curve of the prediction errors remains flat during the foraging patches in Figure 1B. However, the biological signal may have less leeway for a negative than a positive inflection, resulting in a small put positive average. These oscillations of the dopamine signals have been observed (Kobayashi and Schultz 2008; Fiorillo et al. 2008), albeit at longer intervals between targets and after creating obvious expectations of the temporal occurrence of targets by first presenting them regularly. In the present study, participants experienced the irregular appearance of the targets from the outset, and may have incorporated the minor irregularity of the occurrence of the targets in their expectations. Experiment 3 addressed this issue by presenting the targets at regular intervals, thus excluding that the activation in the foraging patches may be due to prediction error generated by delays or anticipations of targets.

Yet another issue is that our model did not represent uncertainty concerning the duration of the foraging patches. As it often happens in reinforcement learning models, the structure of the possible states in the task is provided by the modeller, leaving to the algorithm the assignment of reward level to the states. For this reason, prediction error may have been generated in vivo toward the end of the foraging patch, in contrast to the model which relied on prior knowledge on the possible state sequence. For this reason, as detailed in the Results section, supplementary statistical analyses in Experiment 1 were conducted to show that there were no increasing effects of reward levels at the end relative to the beginning of the foraging patch in VTA/SN.

#### 2.2.2 Simulations

Trials were modelled by discrete steps representing the appearance of the cue, the interval, and the targets in the foraging patch as a finite state approximation to the continuous state. After training the model using eq. (1), we ran the model for 200 trials recording the values taken by *δ*(*s*_*t*_) and *r*_*t*_ at each step. To reflect the characteristics of the BOLD signal generated by these values, Figure 1B displays smooth curves fitted to these values, averaged over the trials. The vertical bars exemplify the values taken by *δ*(*s*_*t*_) and *r*_*t*_ in one exemplary trial. To model the irregular temporal occurrence of the target dots in the foraging patch, we interspersed target (state steps in the model in which a target dot appeared) and non-target episodes (state step with no target dot and no reward) as a finite state approximation to the continuous case. Target and non-target episodes occurred at random between the target episodes. This modelling choice considers the possibility of dopamine firing being sensitive to varying target delays (Experiments 1 and 2; see also Pagnoni et al. 2002; Kobayashi and Schultz 2008; Fiorillo et al. 2008) and is responsible for the signal oscillations at target delays shown in Figure 1B. Smoothed over time and averaged, these oscillations compensate each other at the baseline level. When the targets are presented at regular intervals (Experiment 3), the model simplifies to the quantities *δ*(*s*_*t*_) being uniformly zero during the foraging patches, giving the same modelling average outcome.

Simulations were coded in the publicly available numerical computing language Julia (available from http://julialang.org/; see Bezanson et al. 2017). Figure 1B was drawn with the Julia package Gadfly by Daniel C. Jones (http://gadflyjl.org).

### 2.3 Imaging techniques and data preprocessing

Data were collected with a Prisma 3T Siemens Scanner using a T2*-sensitive echo planar imaging sequence. The acquisition parameters of Experiment 1 were: TR/TE 2460/30 msec, flip angle 82°, 64×64 voxels, FOV 24 cm, 39 2.5 mm slices with a gap of 0.5 mm acquired in ascending order, giving a voxel size of 3×3×3mm. The acquisition parameters of Experiments 2 and 3 were: TR/TE: 2850/40 ms, flip angle 90°, 94×94 voxels, FOV 19.2 cm, 40 2 mm slices with a gap of 0.5 mm acquired in ascending order, giving a voxel size of 2×2×2.5 mm. A 64-channels head coil was used with foam padding to minimize head motion.

After realignment, data were registered to MNI space using the unified segmentation-normalization procedure provided by the SPM package (version 12, Welcome Trust Centre for Neuroimaging, University College London, http://www.fil.ion.ucl.ac.uk/spm/). This algorithm has been shown to be effective in segmenting substantia nigra in EPI volumes (Düzel et al. 2015). In Experiment 1, default settings were used for registration. In the second and third high-resolution experiments we segmented the substantia nigra, red nucleus, and pallidum explicitly as part of the unified segmentation-normalization procedure in SPM12.

The possibility of segmenting the substantia nigra depends on higher concentrations of non-haeme iron in the pars reticulata, which affect the T2*-weighted signal of EPI images by reducing its intensity (Drayer et al. 1986; Vymazal et al. 1995). Similarly high iron concentrations are present in the red nucleus and the pallidum, giving to these brain structures similarly lower signal intensity. We therefore programmed the segmentation algorithm to identify a separate tissue class for these brain structures. Since this segmentation algorithm relies on so-called ‘spatial priors’ to identify the tissue classes, we supplemented the spatial prior maps delivered with the software package SPM12 with priors for the substantia nigra, red nucleus, and pallidum. The priors were created by manually segmenting these nuclei in the Montreal Neurological Institute T1-weighted and T2-weighted images of the ICBM atlas (Fonov et al. 2011), after registering them to the priors provided by the SPM package. The manual segmentation was done with the freely available software ITK-snap (Yushkevich et al. 2006). The segments were smoothed with an isotropic kernel FWHM 1mm before inclusion in the spatial prior set. To accommodate the new priors, existing prior values in the affected voxels were reduced proportionally so as to preserve the unitary sum of the priors. To model the signal densities, we used a parametric approach in which the intensity of the tissue segments was modelled as mixture of Gaussians (a strategy commonly used in structural imaging). We explored several parametrizations of the Gaussians, starting with two Gaussians per tissue class, with the exception of the substantia nigra, red nucleus, and pallidum segments, which were modelled by one Gaussian. To evaluate the success of the combined segmentation-normalization procedure, we computed the maps of the segmented tissues in each participant (by assigning each voxel to the tissue class for which the posterior probability estimated by the segmentation algorithm was largest), and computed maps of the frequency of assignment to the tissue classes after applying the registration parameters. It turned out that the brainstem segmentation was robust to the parametrization of the other tissue classes, and gave very similar results in the final statistical analyses. We chose the final parametrization to best reflect the observed signal density in the rest of the image while simplifying the density models, settling on one Gaussian for all tissues except CSF, bone, and background air, which were each modelled by two Gaussians. As one can see in Figure 2A and 3 below, there were several voxels in which the classification rate of the hypointense region of the substantia nigra reached 100%, showing that in all participants this tissue class was detected by the segmentation algorithm and registered accordingly. This means that there were voxels in all individuals that were classified as substantia nigra and that there was an overlap of these voxels that included all individuals. The segmentations of the red nucleus and pallidum were similarly sensitive. The rate of classification outside the substantia nigra and other nuclei was about zero, showing that the segmentation was selective for the targeted tissue classes.

**Figure 2.**
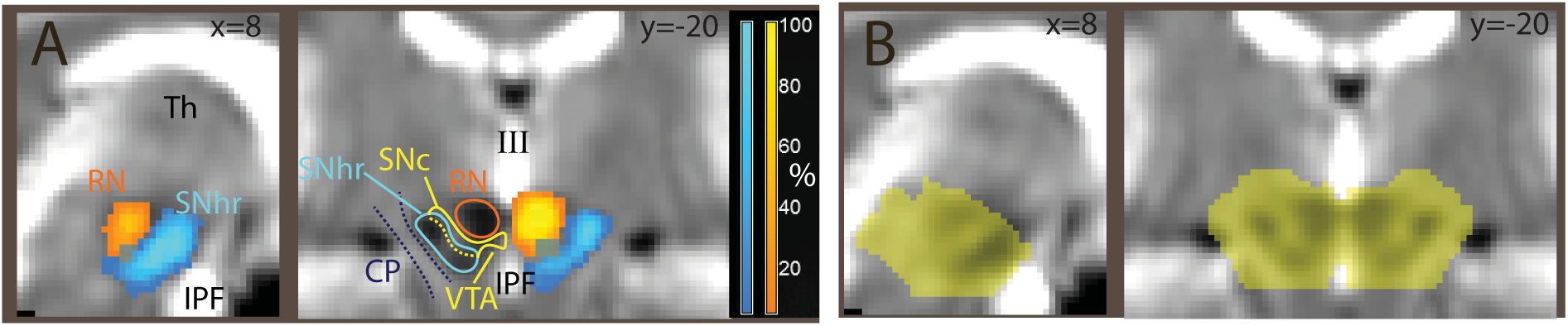
A: Segmentation of the substantia nigra and red nucleus in the second experiment, overlaid on the average EPI image. In blue, tissue classified as substantia nigra by SPM’s segmentation algorithm through its low intensity in the T2*-weighted EPI images used in the segmentation. In orange/yellow, voxels classified as red nucleus. Coordinates in MNI space. B: midbrain mask used for the region of interest correction in Experiments 2 and 3. RN: red nucleus; SNhr: hypointense region, corresponding to iron-rich tissue attributed to the substantia nigra pars reticulata; SNc: region between RN and SNhr not classified as hypointense, attributed to substantia nigra pars compacta; CP: cerebellar peduncle; VTA: ventral tegmental area; IPF: interpeduncular fossa; III: third ventricle.

The approach to segmentation used in the present study differs from those often used in the structural imaging literature concerned with the assessment of the substantia nigra in that we used low-resolution EPI images as input. However, the source of contrast is the same, i.e. the effects of iron on the T2* contrast in MRI images. An argument for the approach presented here is that it is informative of the location of areas affected by this contrast in the modality in which the functional signal originated and of the extent to which these areas were registered together in those volumes. Figure 3 also shows the extent to which the registration was aligned with template images from the ICBM atlas (Fonov et al. 2011). Here, one can see the areas classified as hypointense in all individuals (in white) and the extent of their overlap with the hypointense region in T1-weighted images and its corresponding regions in proton-weighted images. As noted by Düzel et al. (2015), in man most dopaminergic neurons are found in the compacta, which partially overlaps with the reticulata and occupies the region between this latter and the red nucleus.

**Figure 3.**
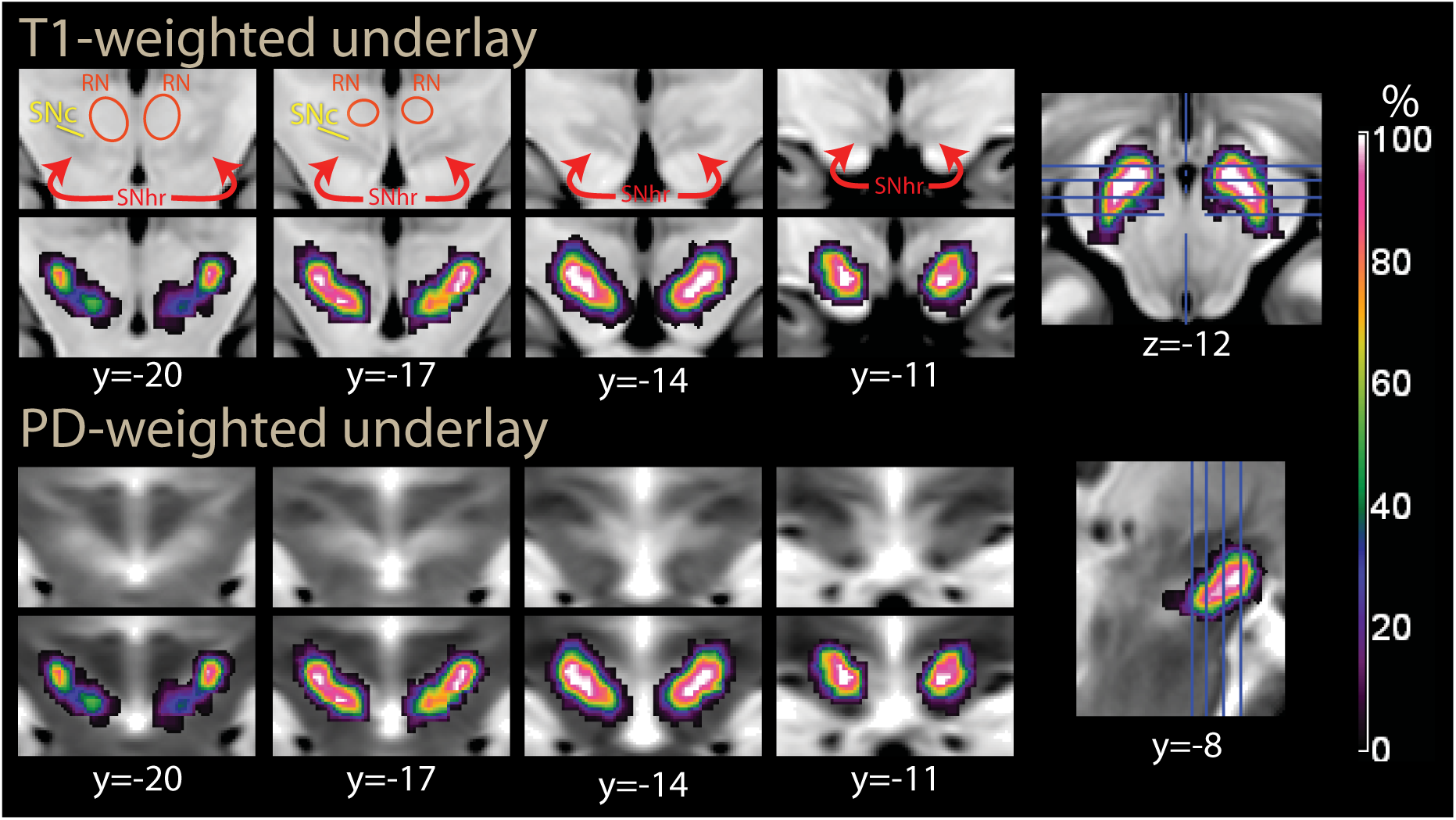
Segmentation of the substantia nigra and red nucleus in the second experiment, as in Figure 2A, overlaid on T1-weighted (top) and proton density-weighted (bottom row) images from the ICBM atlas. In T1-weighted images, the iron-rich region of the substantia nigra is slightly hypointense (SNhr). In proton density-weighted images, it is enclosed in a slightly hyperintense area. The color scale refers to the percent of individuals in which the voxel was classified as belonging to the hypointense region in the T2*-weighted EPI images. MNI coordinates, abbreviations as in Figure 2A.

In registering the data to standard MNI coordinate space, a resampling size of 2mm (Experiment 1) and 1mm (Experiments 2 and 3) was used. Prior to statistical modelling, data were smoothed with a Gaussian kernel of full-width half maximum (FWHM) 8mm (Experiment 1) and 6mm (Experiments 2 and 3). In the model of the interaction between experiments 1 and 2, resampling and smoothing were as in Experiment 1. The smaller smoothing of experiments 2 and 3 was chosen to make use of the increased resolution of the EPI images in these experiments, reflecting prior knowledge from experiment 1 about an effect of interest in the brainstem

### 2.4 Experimental Design and Statistical Analysis

The task consisted of 12 trials, each consisting of a cue displayed for 2 sec., an interval of 3 sec. during which a fixation cross was displayed, and a foraging patch of about 14.5 sec. during which large target dots appeared at irregular intervals in the first and second experiments (according to an exponential schedule bounded between 300 and 1800 msec, and an average interval of about 1220 msec) on the right or on the left of the fixation screen (Figure 1A). Given the focus of the present study on the effects of reward in the foraging patch, the duration of this latter allows modelling it as a block, thus optimizing sensitivity of the design with respect of the dominant frequencies of the BOLD response (Birn et al. 2002). In the third experiment, the same targets appeared at regular intervals of 1250 msec. Participants were required to click on the right or on the left at the appearance of the target dots to collect rewards. The target dots remained on screen for a maximal duration of 800 msec, after which the award would not be collected. Trials were separated by pauses of 10 sec. during which a fixation cross was shown, for a total duration of 6 min for the whole task. Intervals between the presentations of the target dots during the foraging patch were clearly distinguishable from other intervals occurring during the experiment through the continued display of the running total of collected reward. Trials were of two types, depending on the amount rewarded at each correct response (1 cent or 20 cents). Cues and foraging phases belonging in different trial types were identifiable by color. In the main model, we relied on the low frequencies that characterize the BOLD response to smooth a signal approximating reward rates over time in the foraging patches. We presented the probes for the collection of the rewards at a relatively high frequency, modelling the whole patch as a block in the statistical analysis of the neuroimaging data. For administrative reasons we had to change the compensation at the end of the session between the experiments. In the first experiment, participants collected two movie theatre coupons if they had collected at least 20 Euro by the end of the experiment. In the second and third experiments, they received the sum they had collected if they had reached the same threshold. Participants were conditioned on the cue color in a practice run prior to being positioned in the scanner. The three experiments were conducted in three different sessions.

Statistical modelling of neuroimaging data was conducted with the freely available software package SPM12 (www.fil.ion.ucl.ac.uk/spm) using the default settings provided by that package, complemented by the following specifications. The cue and foraging phases of trials were modelled by separate boxcar functions convolved with a standard haemodynamic response of equal amplitude at the cues and foraging phases, and fitted with an intercept term to the MRI data voxelwise using a first-order autoregressive model to account for the temporal correlation of residuals. Contrasts were computed between high and low reward trials for the cue and foraging conditions at the first level. Estimates of contrast coefficients computed in each participant were brought to the second level to model random effects of subjects.

Significance levels were computed with a permutation method (6000 resamples). Clusters were defined by the uncorrected threshold *p* < 0.001 (in corrections for the whole volume) and *p* < 0.01 (in corrections for regions of interest). In the text, the size of cluster *k* is denoted in voxels. The regressor of interest was permuted randomly, and the largest *t* value across voxels (for peak-level correction) or the size of the largest cluster (for cluster-level correction) in the *t* map obtained in the regression was noted in each of 6000 permutation samples (Holmes et al. 1996). In region of interest analyses, the largest value referred to the confines of a region in a predefined anatomical mask. Significance levels for cluster sizes were given by the quantiles of these maximal cluster values. In tables, we also report significance levels at the peak level computed with the random field theory approach. Peak-level corrections (also known as voxel-level) allow for strong control of the family-wise error rate.

The inferential procedure differed in experiment 1 and in experiments 2 and 3. In experiment 1, it was possible to assume a priori an effect in the nucleus accumbens/ventral striatum at the appearance of the cue based on the existing literature. We therefore carried out inference aided by an anatomical region of interest for this region, defined as a box at MNI coordinates x: –12-+12, y: 0-+12; z: –12-+6 (5184 mm^3^), based on the data reported by Abler et al. (2006). In contrast, because no a priori hypothesis was made for the effect of reward in the foraging patches, we conducted the analysis of the respective contrast in the whole volume. In experiment 1 we also used standard native resolution, resampling, registration and smoothing parameters. Analysis of the foraging patch in experiment 1 also aimed at describing cortical effects. In experiments 2 and 3, native resolution, resampling, registration, and smoothing were tailored to detect effects in smaller regions in the brainstem (as detailed above), based on the findings of experiment 1. In experiments 2 and 3, inference was accordingly aided by a region of interest covering the substantia nigra and the surrounding tissue, given by a mask isolating the midbrain created with the freely available software Itk-Snap (Yushkevich et al. 2006; 12461 mm^3^, Figure 2B).

As explained in the Results section, several confirmatory additional models were considered in Experiment 1. To test for a linear trend at the cue, a ‘parametric modulation’ of the convolved haemodynamic response of the cues with trial number was added to the model for the two types of cues separately, and brought to the second level as a contrast between reward levels (interaction reward level x time). To test for variations over time within the foraging patches, they were modelled as a series of events, one for each presentation of the target, and parametric modulations of the resulting convolved regressors were considered. In one model, time within each foraging patch was modelled up to the fourth degree polynomial expansion to assess the course of the signal within this patch. The expansions were orthogonalized by the SPM12 package to those of lower degree. The fit from the main effect and the polynomials expansions were used in the plots of the fitted course of the signal of Figure 6B. The fitted cue signal was computed in the same way from the cue regressors. The contrast of the linear trend for low and high reward trials was brought to the second level to test for systematic increases in the contrast during the progression of the foraging patch. An alternative more flexible model in which the first and last quartiles of the foraging patches (modelled by an appropriate parametric modulator) modelled changes during the foraging patches was also considered for the same purpose.

Plots of the fitted BOLD course of the signal display mean curve and 90% confidence intervals (Figure 6B). These intervals were generated point-wise from the individual BOLD curves fitted at the first level, computing the variance of the signal around the mean curve analogously to a second-level analysis of the model coefficients.

Statistical analyses of behavioural data were conducted with the function *lmer* in the freely available packages lme4 (v 1.0, Bates et al. 2015) and lmerTest (v. 2.0, Kuznetsova et al. 2016) in R (www.r-project.org/). Logistic and linear regression models (for hits and reaction times, respectively) included random effects of intercept and reward levels in subjects to account for repeated measurements. Significance levels in the linear regression were computed using a Satterthwaite approximation for the degrees of freedom. Overlays were composed with the freely available programme MriCroN (https://people.cas.sc.edu/rorden/mricron/index.html).

Overlays were composed with the freely available programme MriCroN (https://people.cas.sc.edu/rorden/mricron/index.html), and annotated with Adobe Illustrator to produce the figures. The template images in Montreal Neurological Institute space are those of the ICBM atlas (Fonov et al. 2011).

### 2.5 Recruitment

Participants were recruited through local announcements on the premises of the University of Ulm. Participants gave their written informed consent. The study was approved by the Ethical Committee of the University of Ulm and was carried out in accordance with the relevant guidelines and regulations (Declaration of Helsinki and Personal Data Protection Convention of the Council of Europe). For Experiment 1, 35 volunteers were recruited (18 females, mean age 23.6±5.1). Descriptive summary statistics of participants in Experiment 2 and 3 were N = 17, 15 females, mean age 22.3±3.2, and N = 20, 14 females, mean age 25.6±4.8, respectively.

## 3 Results

### 3.1 Experiment 1 (irregular alternation of high and low reward trials)

The behavioral data analysis revealed that 29 of the 35 participants (83%) managed to reach the threshold of 20 units of the local currency to win the coupon at the end of the task (mean win: 20.2, std. dev. 0.19). The majority of participants (26, 74%) could catch all targets; the maximal number of failures was four missed targets in one participant. Failure to catch targets was not associated with reward levels in the foraging patches (logistic regression, *t* = 0.2, n.s.). Reaction times at target detection averaged 390.9 msec (std. dev. 72.1), and were 5.6 msec longer in the foraging patches with low reward levels. Notwithstanding its modest size (about 1.4% of the average reaction time), which may not be expected to affect the bold response, this increase was significant (*t* = 3.22, *p* = 0.001). Reaction times also significantly decreased with the progression of the trials (*t* = –9.57, *p* < 0.001) and of targets within trials (*t* = –13.22, *p* < 0.001), presumably as an effect of practice.

In the functional imaging analysis, we computed contrasts of the high vs. low reward conditions at the cue and during the foraging phases separately. At the cue, this contrast activated the ventral striatum bilaterally (Table 1, cluster #4, and Figure 4, top row, blue ovals). Other brain regions activated by this contrast were the left precentral gyrus and the supplementary motor area/middle cingular cortex (Table 1). At lower signal intensity, this signal also activated the basal forebrain and the brainstem in the perirubral area.

**Table 1.**
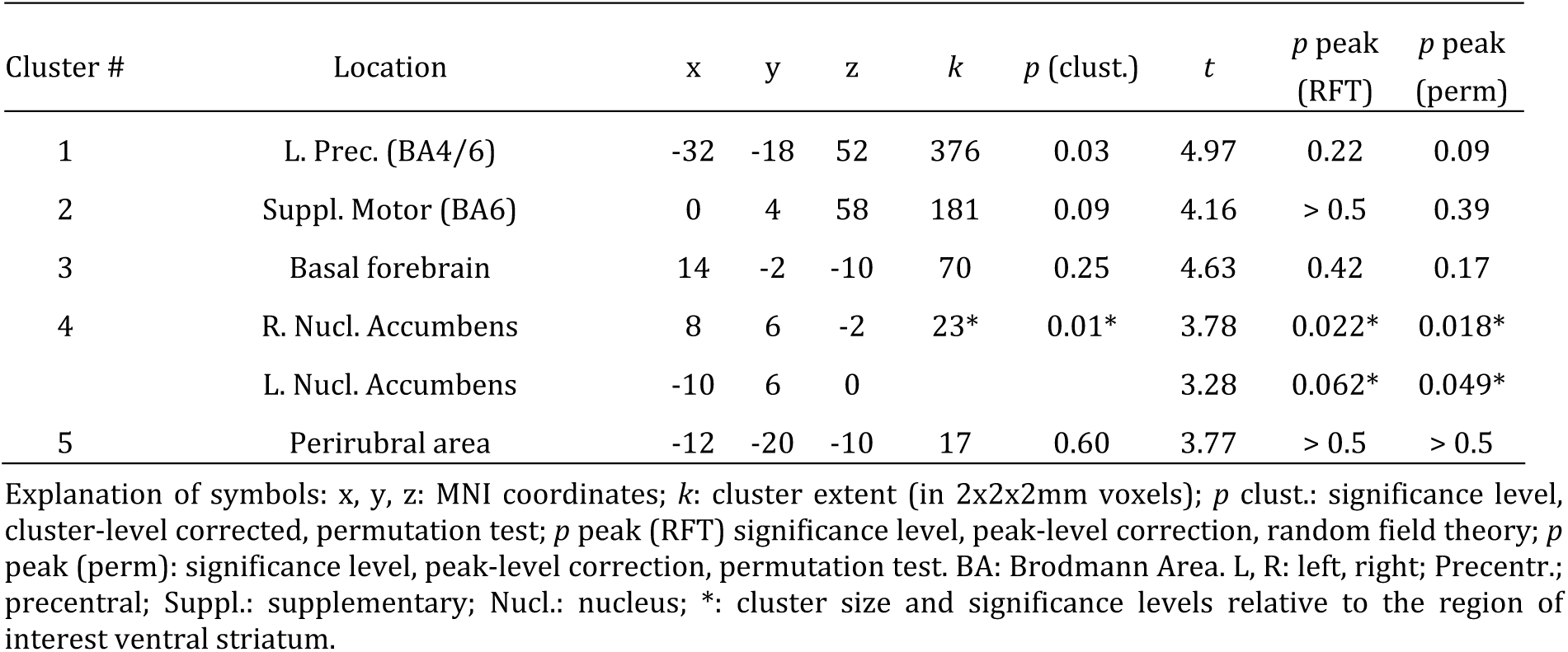
Cue phase, high vs. low reward trials.

| Cluster # | Location | x | y | z | k | p (clust.) | t | p peak (RFT) | p peak (perm) |
| --- | --- | --- | --- | --- | --- | --- | --- | --- | --- |
| 1 | L. Prec. (BA4/6) | -32 | -18 | 52 | 376 | 0.03 | 4.97 | 0.22 | 0.09 |
| 2 | Suppl. Motor (BA6) | 0 | 4 | 58 | 181 | 0.09 | 4.16 | > 0.5 | 0.39 |
| 3 | Basal forebrain | 14 | -2 | -10 | 70 | 0.25 | 4.63 | 0.42 | 0.17 |
| 4 | R. Nucl. Accumbens | 8 | 6 | -2 | 23* | 0.01* | 3.78 | 0.022* | 0.018* |
|  | L. Nucl. Accumbens | -10 | 6 | 0 |  |  | 3.28 | 0.062* | 0.049* |
| 5 | Perirubral area | -12 | -20 | -10 | 17 | 0.60 | 3.77 | > 0.5 | > 0.5 |

Explanation of symbols: x, y, z: MNI coordinates; k: cluster extent (in 2x2x2mm voxels); p clust.: significance level, cluster-level corrected, permutation test; p peak (RFT) significance level, peak-level correction, random field theory; p peak (perm): significance level, peak-level correction, permutation test. BA: Brodmann Area. L, R: left, right; Precentr.: precentral; Suppl.: supplementary; Nucl.: nucleus; \*: cluster size and significance levels relative to the region of interest ventral striatum.

**Figure 4.**
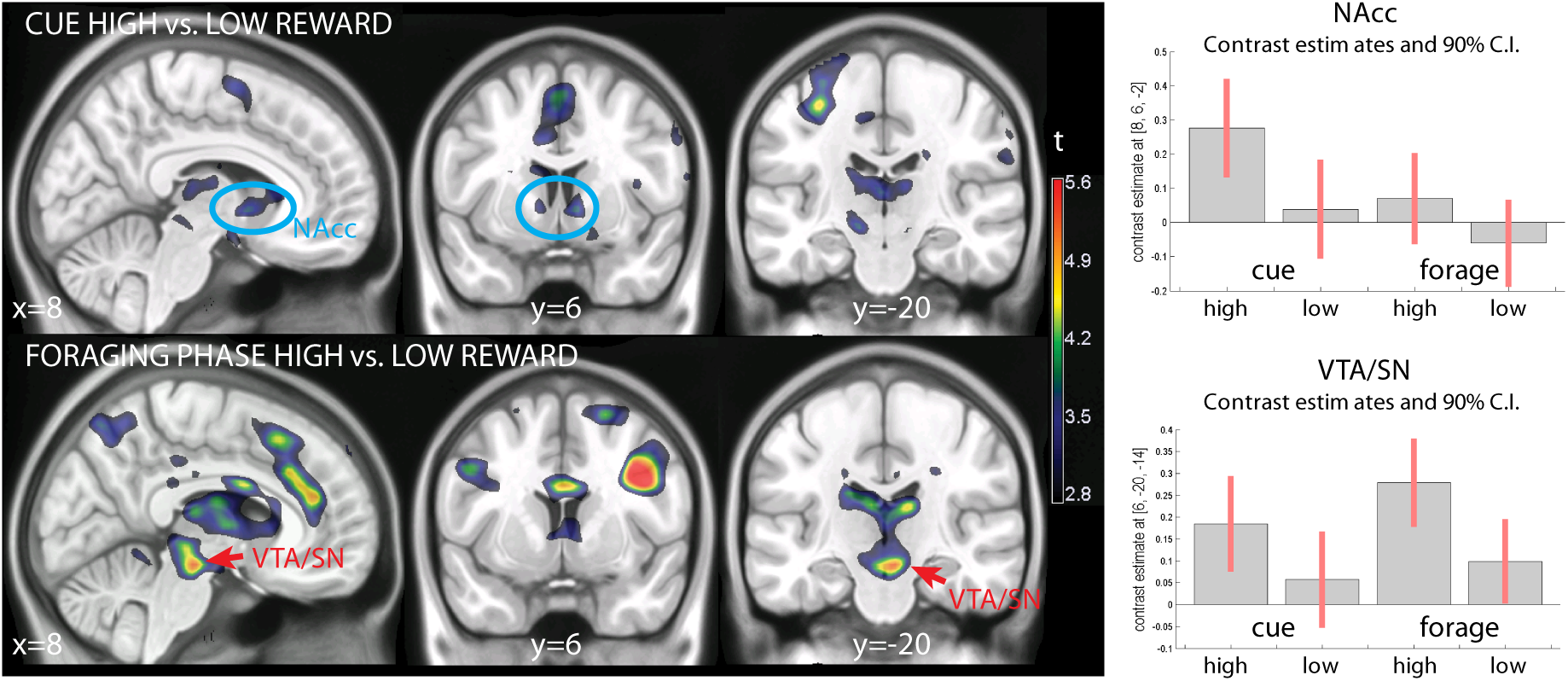
Experiment 1 (N=35, unpredictable alternation of low and high reward trials): high vs. low reward contrast at cue presentation (top row) and during the foraging patches (bottom row). The blue ovals in the top row show striatal activation commented on the text. The red arrows in the bottom row point to activations in the dopaminergic brainstem. On the right, contrast estimates and relative confidence intervals in the nucleus accumbens and ventral tegmental area. Colors show *t* values for the contrast of interest, thresholded for illustration purposes at *p* < 0.005, uncorrected, overlaid on a template brain. NAcc: nucleus accumbens; VTA: ventral tegmental area; C.I.: confidence intervals.

The contrast of different reward rates during the foraging patches aimed at detecting areas responding to mean reward rates. This contrast activated the brainstem in a region encompassing the ventral tegmental areas/substantia nigra (VTA/SN, x, y, z: 6, –20, –14, *t* = 5.44, *p* = 0.024, peak-level corrected for the whole brain; Figure 4, bottom row, red arrows, and Table 2, cluster 5). This cluster of activation extended posteriorly into the deep collicular tissue at the border with the pons and into the superior colliculus (Table 2, cluster 5).

**Table 2.**
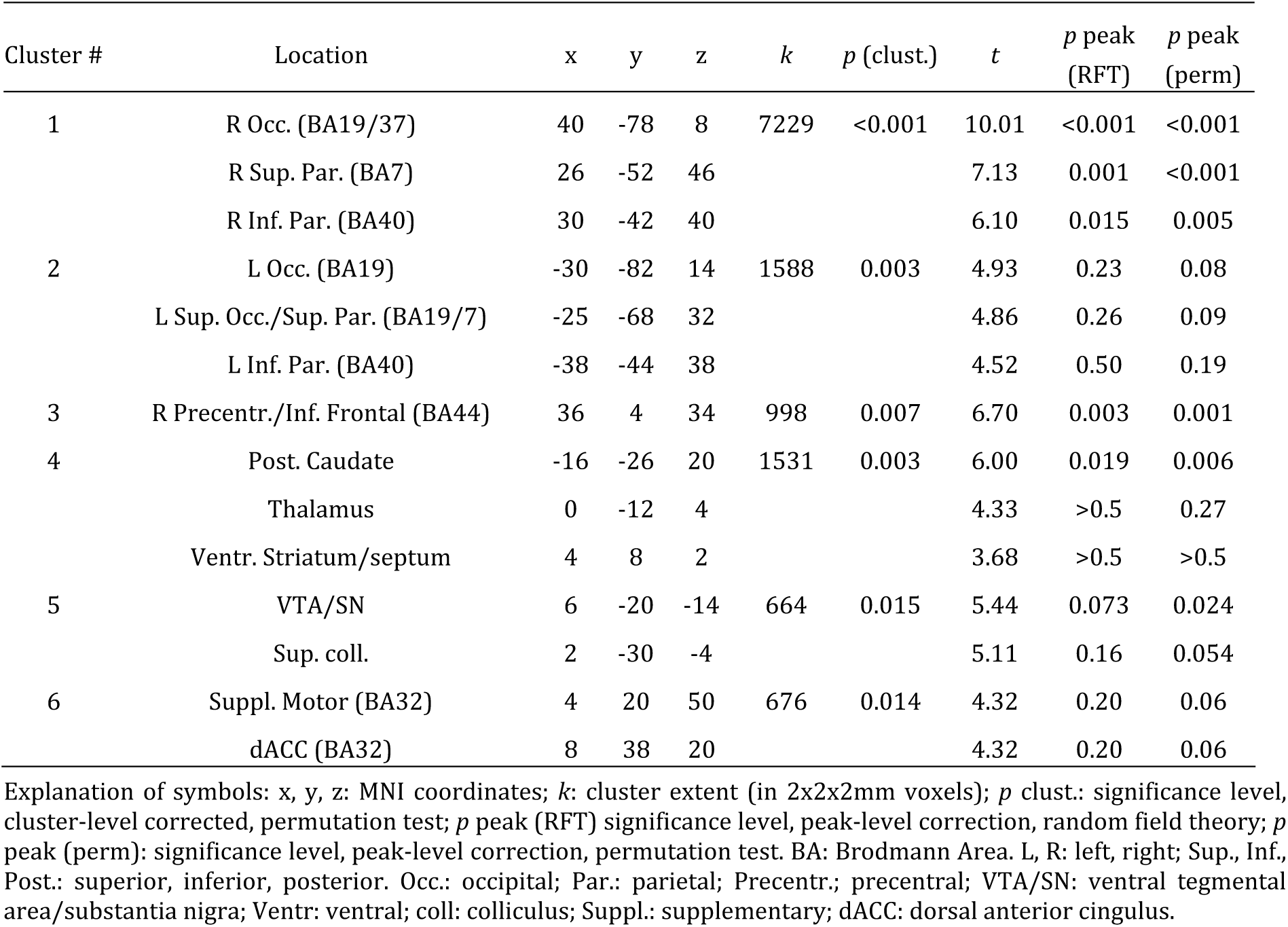
Foraging phase, high vs. low reward trials.

| Cluster # | Location | x | y | z | k | p (clust.) | t | p peak (RFT) | p peak (perm) |
| --- | --- | --- | --- | --- | --- | --- | --- | --- | --- |
| 1 | R Occ. (BA19/37) | 40 | -78 | 8 | 7229 | <0.001 | 10.01 | <0.001 | <0.001 |
|  | R Sup. Par. (BA7) | 26 | -52 | 46 |  |  | 7.13 | 0.001 | <0.001 |
|  | R Inf. Par. (BA40) | 30 | -42 | 40 |  |  | 6.10 | 0.015 | 0.005 |
| 2 | L Occ. (BA19) | -30 | -82 | 14 | 1588 | 0.003 | 4.93 | 0.23 | 0.08 |
|  | L Sup. Occ./Sup. Par. (BA19/7) | -25 | -68 | 32 |  |  | 4.86 | 0.26 | 0.09 |
|  | L Inf. Par. (BA40) | -38 | -44 | 38 |  |  | 4.52 | 0.50 | 0.19 |
| 3 | R Precentr./Inf. Frontal (BA44) | 36 | 4 | 34 | 998 | 0.007 | 6.70 | 0.003 | 0.001 |
| 4 | Post. Caudate | -16 | -26 | 20 | 1531 | 0.003 | 6.00 | 0.019 | 0.006 |
|  | Thalamus | 0 | -12 | 4 |  |  | 4.33 | >0.5 | 0.27 |
|  | Ventr. Striatum/septum | 4 | 8 | 2 |  |  | 3.68 | >0.5 | >0.5 |
| 5 | VTA/SN | 6 | -20 | -14 | 664 | 0.015 | 5.44 | 0.073 | 0.024 |
|  | Sup. coll. | 2 | -30 | -4 |  |  | 5.11 | 0.16 | 0.054 |
| 6 | Suppl. Motor (BA32) | 4 | 20 | 50 | 676 | 0.014 | 4.32 | 0.20 | 0.06 |
|  | dACC (BA32) | 8 | 38 | 20 |  |  | 4.32 | 0.20 | 0.06 |

Explanation of symbols: x, y, z: MNI coordinates; k: cluster extent (in 2x2x2mm voxels); p clust.: significance level, cluster-level corrected, permutation test; p peak (RFT) significance level, peak-level correction, random field theory; p peak (perm): significance level, peak-level correction, permutation test. BA: Brodmann Area. L, R: left, right; Sup., Inf., Post.: superior, inferior, posterior. Occ.: occipital; Par.: parietal; Precentr.: precentral; VTA/SN: ventral tegmental area/substantia nigra; Ventr.: ventral; coll.: colliculus; Suppl.: supplementary; dACC: dorsal anterior cingulus.

The coefficients estimates of Figure 4 also show that there was some differential activation of this area during the presentation of the cue, which however failed to reach any level of significance. Coming back to the foraging phase, also the caudatus was significantly active (Table 2, cluster 4). The ventral striatum was activated by high reward rates in the foraging patches together with a more posterior are extending into the medial thalamus, without however reaching significance after correction. In the cortex, a predominantly right-lateralized visual and attentional network was active (Table 2, clusters 1-3 and 6).

To verify the local distribution of the signal in the foraging phase of the experiment, we repeated the analysis after smoothing the data with a smaller 6mm kernel. This analysis revealed a predominantly right-lateralized effect of reward in the VTA/SN, peaking in the right SN (x, y, z: 6, –20, –14, *t* = 5.77, *p* = 0.021, peak-level corrected, and *k* = 426, *p* = 0.011, cluster-level corrected for the whole brain). In Figure 5, we show this effect overlaid on a proton-weighted template brain, where SN is visible as a brighter band of tissue (Oikawa et al. 2002). Figure 5B shows the effect of reward levels in transversal slices of the mesencephalon.

**Figure 5.**
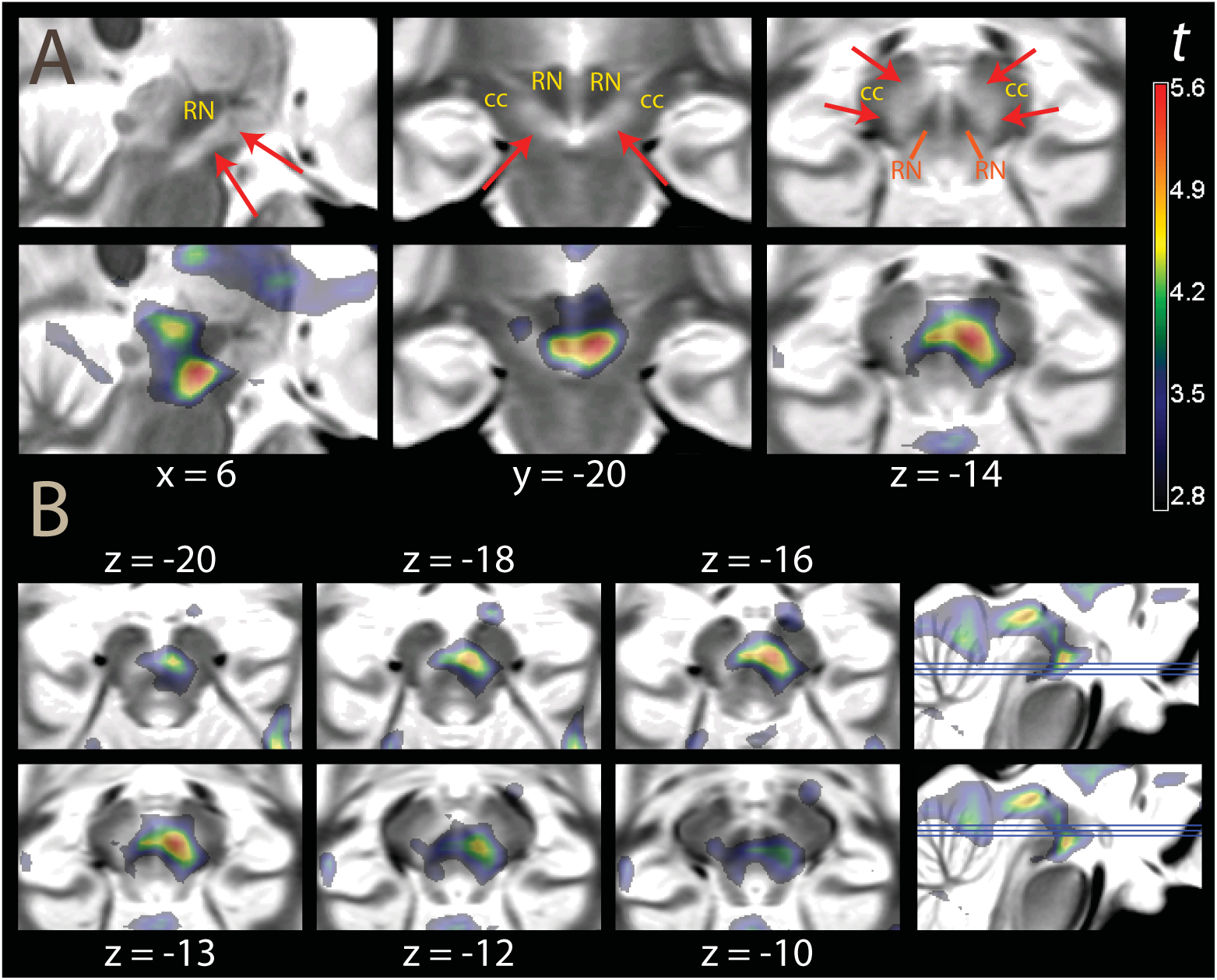
Inset showing the effect of reward levels in the foraging phase (data smoothed FWHM 6mm for improved resolution) on a proton-density weighted template brain. A: statistical maps centered at the pial voxel. In the upper row, the template image without overlay. The SN is visible as a brighter strip of tissue (red arrows), while the crus cerebri (cc) has a darker appearance. Note that here, in contrast to Figure 2A, there is no contrast between the iron-rich portion of SN and the rest. RN: red nucleus. B: transversal slices. Montreal Neurological Institute coordinates. The thresholding for visualization and the color scale are the same as in Figure 4.

**Figure 6.**
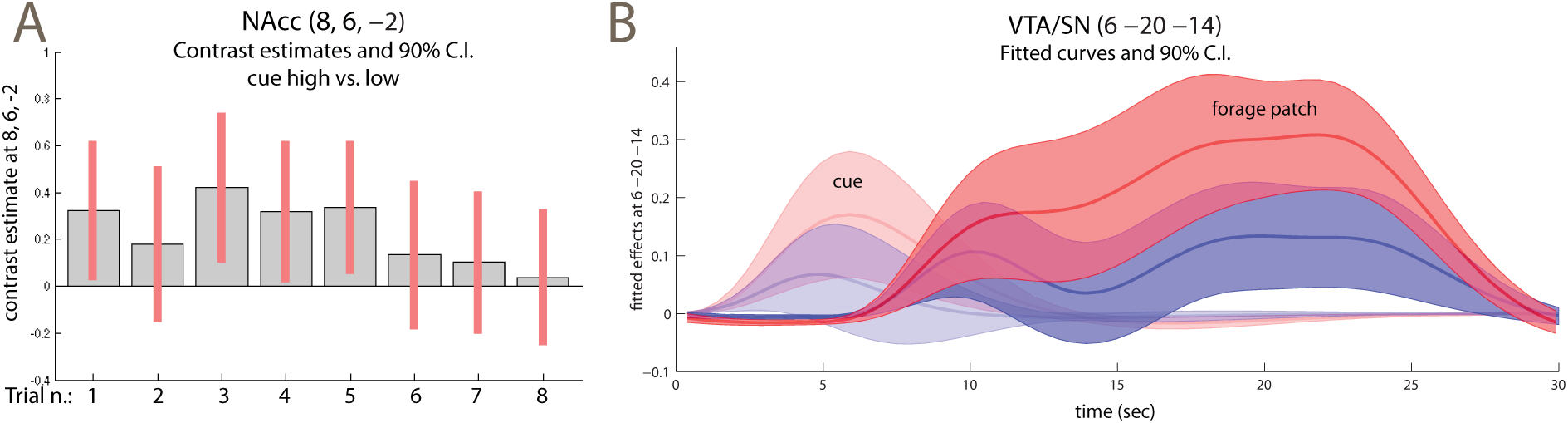
A: contrast estimates high vs. low reward cue computed separately over the trials of the experiment. B: time course of the fit and 90% confidence intervals of the cue and foraging patch at VTA/SN location (the fitted signal of the overall model, at the net of the intercept, is given by the sum of the effects of cue and foraging patch). The time course for the average high and low reward trials are in red and blue, respectively. The signal course of the forage patch was obtained by fitting a fourth-degree polynomial on the events given by the appearance of the targets, thus revealing the course of the signal in the patch. The signal shows a tendency to rise during the patch, but is otherwise fairly uniform. Like the contrast estimates of the bottom row of Figure 4, the time course of the fitted cue signal suggests that low reward levels were activating VTA/SN relative to fixation much like high levels, even if less intensively.

We conducted supplementary analyses to verify some of the assumptions intrinsic in the simple model of reward processing of Figure 1B. First, this model assumed that conditioning to the cues had already taken place due to the practice given to participants prior to the scanning session. This justified not modelling learn effects during the scanning session. To verify this assumption, we modelled the cues in the eight trials separately, and computed the respective contrast estimates (Figure 6A). If participants had not been conditioned to the cue at the beginning of the experiment, we should observe no or a lower effect of reward levels in the ventro-striatal region in the first few trials. As the Figure shows, however, this was not the case; on the contrary, the amplitude of the reward effect decreased over time (a model that included the time trend over the trials gave non-significant negative coefficients).

Second, both the regressor used to assess the effects of reward in the foraging patches and the computational model tended to impose uniformity of effects in the foraging patch. In the case of the regressor, this was due to modelling the foraging patch as a homogeneous block reflecting the constant level of expected reward. In the case of the computational model, this is because it used prior knowledge of the length of the foraging patches, so that learning only consisted of identifying the reward levels of cue and foraging patches. This neglects possible learning effects about the length of the patch. Uncertainty about the opportunity for reward might increase toward the end of the foraging patch, as the cue recedes in time. To address the adequacy of the model in this respect, we created an alternative BOLD regressor for the foraging patches from a series of short events of equal duration (for each target presentation), and computed time trends up to the fourth-degree polynomial to display the course of the data. The plot of the fitted polynomial curves (Figure 6B) shows similar courses in the low and high reward conditions, with the high reward trials always showing more activity than the low reward trials, albeit less so at the beginning and at the end of the patch. We looked at the effect of reward levels on linear trends to verify that this effect remained approximately constant during the foraging patch. This trend was positive, but did not reach significance (6, –20, –14, *t* = 1.47). We also assessed the trend in an alternative model where we contrasted the last with the first quartile of the foraging patch, obtaining similar findings (*t* = 1.67). As whole, Figure 6B shows that the model of sustained activity in the foraging patches reflecting levels of reward was fairly adequate in this region.

### 3.2 Experiment 2 (regular alternation of high and low reward trials)

In Experiment 2 (N=17) we regularly alternated high and low reward trials, and informed participants of this fact. According to the temporal difference model, this predictability should affect the learning signal conveyed by the cues, because these latter now fail to provide new information about the impending rewards. This prediction has been verified in previous functional neuroimaging studies, which found reduced or absent ventral striatal activation when the cues are predictable (Schultz et al. 1993; Berns et al. 2001). However, if our model is correct, brain signals tracking mean reward rates should remain unaffected by changes in the predictability of the cues, because they do not depend on the generation of a prediction error. Since in the first experiment we had observed an effect of mean reward rates in the midbrain, we increased the resolution of images in the second experiment and modified the registration algorithm to segment these brainstem structures explicitly, and applied the correction for multiple testing to a mesencephalic region of interest (see the Methods section for details). The explicit segmentation had two purposes: first, since the registration took into account the segmentation output, it may have realigned the substantia nigra more precisely; second, the output of the segmentation provided information on the extent of coregistration of this structure in the final normalization and assists in the localization of functional effects.

The contrast high vs. low reward at the presentation of the cue failed here to detect any effect of reward rates in the ventral striatum (no voxels survived the cluster-definition threshold), in contrast to the results of Experiment 1. The difference between the effects in the two experiments was confirmed by a significant interaction (x, y, z: 4, 0, –4, *t* = 3.63, *p* = 0.02, peak-level corrected, and *k* = 20, *p* = 0.05, cluster-level corrected for the region of interest). To determine significance levels in the contrast assessing activity in the foraging patches, we computed corrections for a region of interest given by the midbrain, as our intent was replication of the data from Experiment 1 in this region. This contrast detected a cluster of activity centered on the VTA/SN and extending posteriorly towards the superior colliculi (–7, –22, –13, *t* = 5.95, *p* = 0.02, peak-level corrected, and *k* = 437, *p* = 0.03, cluster-level corrected for the region of interest, and 9, –20, –10, *t* = 4.62, *p* = 0.10, *k* = 454, *p* = 0.03, same corrections; see Figure 7A) and more caudally towards the posterior pontine border (8, –26, –8, *t* = 3.74, n.s.).

**Figure 7.**
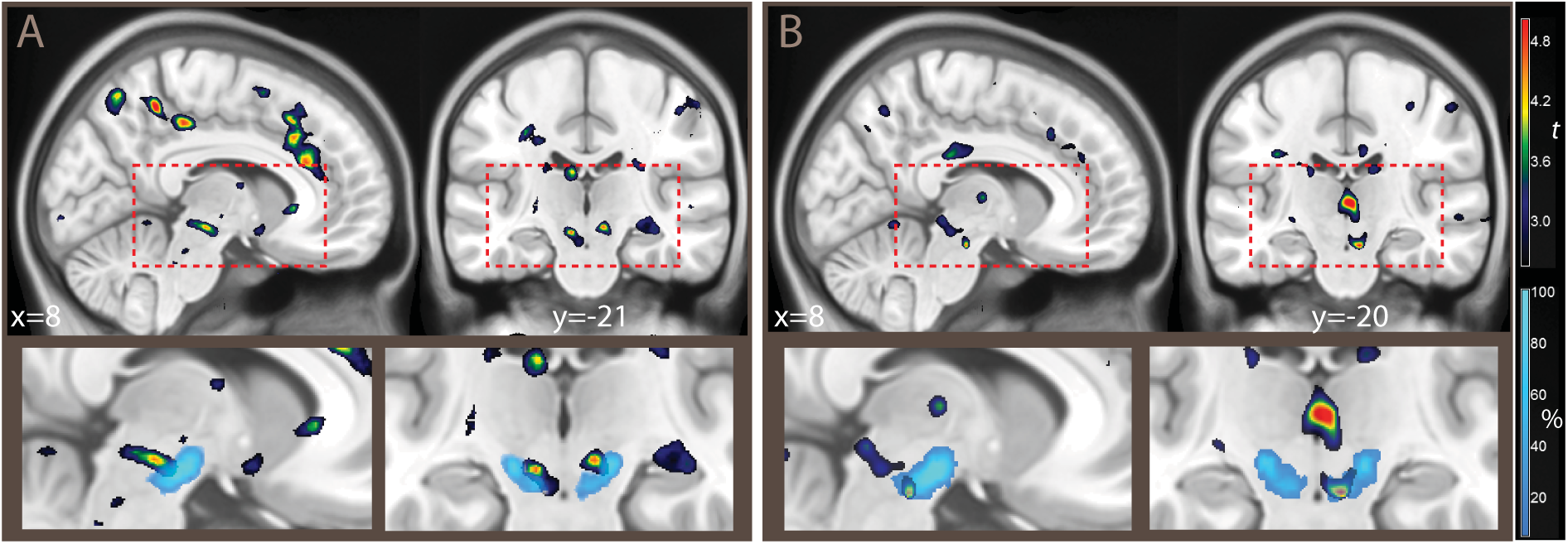
A: Experiment 2 (N=17, regular alternation of high a low reward trials). B: Experiment 3 (N=20, regular occurrence of targets). The overlays show *t* values for the contrast high vs. low reward during the foraging patches, thresholded for illustration purposes at *p* < 0.01, uncorrected. In the insets in the bottom row, percentages of classification of voxels in the SN pars reticulata across the sample are shown by overlays in light blue (from the segmentation algorithm used in the registration). The classification data are the same as in Figures 2A and 3.

### 3.3 Experiment 3 (regular appearance of targets in the foraging patches)

In Experiment 3 (N=20) we increased the predictability of the targets by presenting them at regular intervals. The irregular presentation of the targets corresponds to how opportunities for reward would arise in most realistic circumstances. However, since the temporal difference model computes expectations of reward at each time point, delays or early occurrences of targets perturb the match between expected and observed rewards (Kobayashi and Schultz 2008; Fiorillo et al. 2008). The possible effect of these oscillations is important, because it raises the possibility that prediction error, not the mean reward rates, may be responsible for the motivational signal in the foraging patches. If the signal arising during the foraging patches is distinct from a prediction error signal, Experiment 3 should replicate the finding of the VTA/SN activation, even if the targets were always presented at the same intervals. As in Experiment 2, we increased the resolution of the volumes and performed the analysis in the midbrain region of interest.

Consistently with the replication hypothesis, activation was detected in the right SN (x, y, z: 7, –20, –18, *t* = 5.06, *p* = 0.0475, peak-level corrected for the region of interest of the mesencephalic brainstem; Figure 7B) and, without reaching corrected significance, in the VTA (4, –24, –16, *t* = 3.72). As in the previous experiments, the activation extended posteriorly into the superior colliculus (5, –32, –5, *t* = 5.39, *p* = 0.03, peak-level corrected, same cluster) and into a more caudal region at the posterior boundary with the pons (9, –28, –12, t = 3.25, n.s.; 4, –30, –20, t = 3.58, n.s.). These activations formed a single cluster (*k* = 1429, *p* = 0.001, cluster-level correction for the region of interest).

## 4 Discussion

The novel finding of the present study was the detection of parts of the brainstem active during high-reward foraging patches in conditions in which vigor of responses were matched. The potential significance of this finding is related to the detection of a neural substrate associated with levels of reward and motivation in the absence of the prediction errors conveyed by the cues. The different sensitivity to predictability of rewards in the ventral striatum and this area suggest that the neural substrates activated by high levels of reward during the foraging patches were at least partially dissociable from those activated by high-reward cues. This is consistent with insights coming from the computational model (which distinguishes between levels of rewards and the cues predicting these levels) as well as with intuitive reasoning (which suggests that the motivation to act to exploit perceived opportunities should be present even if the conditions that yield rewards are fully predicted).

The findings concerning the signal associated with the cue replicated the observations of a now fairly extensive body of functional neuroimaging literature. In the first experiment, differences in reward rates at the cue activated the ventral striatum (Berns et al. 2001; McClure et al. 2003b; O’Doherty et al. 2003; Abler et al. 2006; Li et al. 2006; Tobler et al. 2006; Seymour et al. 2007; Rutledge et al. 2010). This activity was present from the first trial, confirming that the cue was predictive from the outset of the experiment. This ventro-striatal activity was consistent with differential reward values being assigned to the trials, even at equal pay-outs. However, in the second experiment, where the cue carried no information on the level of reward of the impeding trial, we did not observe the same activation of the ventral striatum in association with the presentation of the cue. This finding is consistent with previous observations in similar studies that no ventral striatal signal is present at the presentation of cues or reward if their occurrence is predicted (Abler et al. 2006), as when the cues follow a predictable sequence (Berns et al. 2001), and with the prediction of the temporal difference model (Schultz 2015). Together, the findings about the cue characterize the activation in the ventral striatum as the known correlates of prediction errors about available rewards.

In contrast, it is difficult to explain the activation in the brainstem, which was consistent with its localization in the VTA/SN, entirely as a correlate of prediction error. Firstly, the VTA/SN signal in the foraging patch was present also in the second and third experiments, in contrast to the signal at the cue in the ventral striatum, as we would expect from a signal that did not depend on the generation of a prediction error. Secondly, the signal in this area was sustained over the whole foraging patch, suggesting that our simplified model was sufficient to capture activity in this region. Any prediction error coding in the foraging patch, not countenanced by the simplified model adopted here, could only have gradually emerged during the foraging patch because of the possible uncertainty about the appearance of new targets as the cue became progressively more distant in time. Under an alternative model in which prediction error accounts for the activity in the foraging patch, we can still assume no prediction error at the beginning of the patch because in this phase the reward level was clearly predicted by the cue (Berns et al. 2001; Abler et al. 2006). Hence, to account for the effect in the VTA/SN in the foraging patch in this model, this late effect should have been large enough to produce a significant effect of reward for the averaged foraging patch as a whole. This, however, was not the case: the time course of the signal in the VTA/SN was similar in the high and low reward trials. There was some activation at the cue, larger in the high reward trials, and the effect of reward tended to rise towards the end of the foraging patch, but these effects were very far from being significant, in contrast to the constant activity in the foraging patch. Hence, this effect was too small to explain the observed difference in the signal associated with reward levels. Finally, the ventral striatum was not significantly active during the foraging patch, further supporting a different origin of the signal observed during this phase in VTA/SN. If the activity fitted by the foraging block regressor had been attributable to prediction error, we would have expected it to be present in the ventral striatum as well. As a whole, these findings suggest that the activations in VTA/SN in the foraging patch did not depend on the lack of predictability of the level of reward of the trials, but are consistent with the hypothesized effect of the level of perceived reward opportunity in itself. The implication of our design, in which motor efforts and pay-outs were matched across reward levels, is that the reported activity refers to automatic processing of reward levels even when they were not relevant for the task at hand. This is consistent with the relatively poor activation in ventro-striatal areas during the foraging patches, in contrast with studies that have investigated effects of reward levels on the vigor of motor response (Pessiglione et al. 2007; Kurniawan et al. 2010).

These findings are relevant to understand how representations of potential rewards motivate action in different phases of the interaction with the environment. The signal associated with the cue, which carries information on the appearance of an impending reward and is modeled by a prediction error, is widely thought to have motivational effects such as the activation of approach or seeking behavior (Montague and Berns 2002; McClure et al. 2003a). Intuitive reasoning, however, suggests that the motivation to act to exploit perceived opportunities should be present even if the conditions that yield rewards are fully predicted. This reasoning is captured by models of dopamine signaling that emphasize the importance of opportunity costs (Niv et al. 2007; Phillips et al. 2007). The present findings provide evidence on the neural substrates associated with these different aspects of the computations relevant to motivate behavior in the presence of rewards.

While several hypotheses may be formulated about the origin of the activation detected during the high-reward foraging patches, they all support the notion of the motivational significance of this signal. According to an influential hypothesis (Niv et al. 2007), the phasic and tonic modalities of dopaminergic release, which arise in the ventral tegmental area under distinct physiological control and influence the dynamics of dopamine release in the nucleus accumbens (Grace 1991; Floresco et al. 2003; Goto and Grace 2005), may respectively correspond to prediction errors and levels of reward. While the present study cannot distinguish between phasic and tonic signals, as is in the nature of functional neuroimaging data, the activation in VTA/SN detected here is consistent with the hypothesis of tonic dopamine release. Recent neurophysiological data have shown that even when the phasic dopamine firing is blocked, ‘non-phasic’ dopamine signaling still occurs, and the capacity of animals to sustain efforts to obtain rewards is preserved (Zweifel et al. 2009). These findings support of the notion of distinct dopamine signals with motivational significance other than the phasic discharge associated with the prediction error.

While fMRI data cannot be conclusive about the specific involvement of dopamine in the brainstem activations detected in the absence of prediction errors, data from neurophysiological studies in laboratory animals provide a fairly large and consistent body of evidence in this respect. Dopaminergic brainstem nuclei are involved in basic visual-attentional processes in coordination with deep collicular structures (Horwitz 2000; Comoli et al. 2005; Redgrave and Gurney 2006; Düzel et al. 2009; Bromberg-Martin et al. 2010). Dopaminergic neurons in VTA/SN have been shown to fire in response to salient visual stimuli early after their presentation, before full recognition of stimuli and before the prediction error can have been computed (Schultz and Romo 1990; Redgrave and Gurney 2006; Nomoto et al. 2010). This early firing has been shown to be modulated by the existence of a rewarding context (Kobayashi and Schultz 2014), suggesting the existence of a sensitisation mechanism predisposing the system to be reactive to stimuli when they are likely to be rewarding (for a review, see Schultz 2015). In studies from the animal learning literature, the association with the rewarding context has been shown to facilitate task responses (Schultz 2016). In our data, high reward levels were associated with a very small but significant shortening of reaction times in target detection. Because in the present experiment the cues set the context for different levels of reward, one may expect this mechanism to be active during the foraging patches. We suggest that the notion of ‘rewarding context’ of this literature, i.e. a situation in which appearing stimuli are likely to lead to rewards, is similar if not identical to the notion of ‘opportunity cost’. The activations in the posterior brainstem detected in the foraging patches in the present study are consistent with recruitment of the deep collicular structures.

During the foraging patches, the effect of reward extended posteriorly from the substantia nigra towards the tegmentum. In the brainstem, an increasing body of evidence implicates the peduncolopontine tegmentum/laterodorsal tegmental nucleus (PPTg/LDTg) in modulating brainstem activity in rewarding contexts (for reviews, see Düzel et al. 2009 and Gut and Winn 2016). PPTg responds at very short latencies, suggesting that it may drive the early firing of VTA/SN (Sesack and Grace 2010). Hence, PPTg activity in concert with VTA/SN is thought to influence visual sensory processes, locking attention onto stimuli from the external environment (Kobayashi and Isa 2002), and prioritizing attentional deployment and quick reaction when the context suggests that external stimuli may be associated with rewards (Gut and Winn 2016).

In the foraging patches, other areas more active in high reward trials were a prevalently right-lateralized fronto-parietal network, which in previous studies was associated in conjunction with PPTg and the medial thalamus with sustained attention tasks (Paus et al. 1997). In the literature on the activation of brainstem nuclei such as PPTg, briefly reviewed above, and in the discussion of the role of quick dopamine firing prior to the computation of prediction error, the emphasis is on the attentional consequences of the sensitization to rewarding contexts (Schultz 2016), due to the importance of resolving uncertainty on the spatial and temporal appearance of stimuli of high reward value quickly.

The possible involvement of the dopamine signal in attention was formulated in parallel to its involvement in prediction error for rewards (Schultz et al. 1997; Servan-Schreiber et al. 1998). More recently, evidence has accumulated on the effect of reward on selective attention and orienting. In visual attentive tasks, it has been shown that the increased detection of reward-cued targets (Anderson et al. 2013) is associated with brainstem activity in man (Hickey and Peelen 2015), presumably representing dopaminergic facilitation. The effect of dopamine may extend to the attentional component of effort. An application of the effortmodel has been proposed within a broader notion of motivation encompassing facilitating effects on sustained attentional efforts (Kurzban et al. 2013). In a behavioural study in which the amount of reward was systematically varied, an effect of levodopa was the reversal of the insensitivity induced by fatigue on the invigorating effect of higher reward rates (Beierholm et al. 2013). Furthermore, it has been noted that data from animal studies that suggest that dopamine may facilitate the capacity to remain on task (Nicola 2010). In a study of dopamine transporter knockout mice, an increase in persistence in hyperdopaminergic mice in pursuing less rewarding options has been observed, rather than effects on movement such as the tendency to run to the levers operating reward delivery (Beeler et al. 2010). Beside the well-known consequences on physical activity in the free operant setting, the invigorating effects of motivational factors arising from opportunity costs might therefore involve also facilitation of cognitive efforts (Kurniawan et al. 2011). This may facilitate sustaining attentional levels in highly rewarding conditions even when the repetitive nature of the task makes it less amenable to the motivational effects of prediction errors.

The investigation of neural substrates activated when the consistency condition is met is a task for future studies in which the foraging or similar paradigms, in combination with formal models, may contribute to the identification of psychopathological states related to functioning in the positive valence/appetitive motivation domain. Given the importance of models of dopaminergic activity and reward in psychopathology, the capacity to distinguish between different phases of appetitive motivational processes is likely to add to our understanding of pathological conditions.

## Acknowledgements

This study was supported through a NEURON-Eranet grant by the German Federal Ministry for Education and Research (BMBF, Project BrainCYP, Grant. No. 01EW1402B) and by collaborative grants with the Federal Institute for Drugs and Medical Devices (BfArM, Bonn, Grants No. V-15981/68502/2014-2017 and V-17568/68502/2017-2020) to RV. A preliminary version of this work was presented at the 22nd Annual Meeting of the Organization for Human Brain Mapping, Geneva June 26-30, 2016, and at the 14th Society for Neuroeconomics Annual Meeting, Berlin August 28-30 2016.

The authors declare no competing interests.

